# A chromosome-level reference genome of a Fabaceae species yam bean (*Pachyrhizus erosus*)

**DOI:** 10.1101/2023.09.26.559645

**Authors:** Fengjiao Bu, Fan Jiang, Caishun Zhang, Lihua Yuan, Wei Fan, Xinyao Xiong

## Abstract

Yam bean (*Pachyrhizus erosus* L.), belonging to the family Fabaceae, is an important but underutilized root crop. Here, we generated a high-quality chromosome-level reference genome of yam bean by PacBio HiFi and Hi-C sequencing, with assembly size of 539.0 Mb, contig N50 of 25.6 Mb, and BUSCO complete rate of 99.3%. Then, we anchored 94.4% of the contig sequences into 11 pseudo-chromosomes, and assembled the telomeres at 86.4% (19/22) of the chromosome-ends. A total of 44,692 protein-coding genes were predicted, with the BUSCO complete rate of 99.3%, comparable to that of the genome assembly. Compared to the previously reported yam bean genome, the current assembly has a 1,388-fold increase in contig N50 size, and 12.2% and 24.3% increase in BUSCO complete rate for the genome sequence and gene set, respectively. Evolutionary analysis revealed that yam bean diverged from the clade of soybean and *Pueraria lobata* var. *montana* 22.5 MYA. This high-quality genome assembly will greatly facilitate the breeding of yam bean based on the genetic and genomic methods.

## Introduction

Yam bean, or jicama, has been cultivated as root crops for formation of edible roots with higher yields. The three most cultivated varieties of yam bean are *Pachyrhizus erosus, P. tuberosus* and *P. ahipa*. The genus is closely related to Glycine max (Fernandez *et al*., 2021). Unlike soybean, it is only regionally cultivated in Southeast Asia, Mexico and few other countries for its juicy tuberous roots which is rich in Vitamin C, protein and minerals. The sweet and crispy whitish roots are suit for cooking or eating raw as salad or fruit. It can also be used for producing flour and drinks. Yam bean can generate large aboveground biomass under warm and humid conditions, which makes it a potential forage crop. As a nitrogen-fixing legume plant, it can elevate soil fertility during crop rotation. Besides, due to higher rotenone and other compounds in aboveground tissues, it is capable of natural insect resistance. It also shows good adaptability and higher abiotic stress tolerance with reliable yield. Based on the above description, yam bean is a highly neglected crop (Adewale and Nnamani, 2022; Council, 2006). However, lacking higher quality genome and other genetic information which in turn undermine its agricultural utilization and improvement.

Here we report a chromosome-level reference genome of *Pachyrhizus erosus* by using PacBio HiFi and Hi-C sequencing. This high-quality reference genome will pave the way for future genetic improvement of yam beans.

## Materials and Methods

### Plant materials and sequencing

The yam bean cv. Fanji was grown in a greenhouse of Agricultural Genomics Institute at Shenzhen, Chinese Academy of Agricultural Sciences, Guangdong province, China. For genomic assembly, the genomic DNA was extracted from young leaves using the Hi-DNAsecure Plant Kit (cat. no. DP350; TIANGEN, China). Then, one PCR-Free DNA library with 350-400 bp inserts was constructed and paired-end sequenced (2 × 150 bp) on an Illumina NovaSeq 6000 system (Illumina, USA), and two libraries with 15-20 kb inserts were prepared and sequenced on a PacBio Sequel II platform using Circular consensus sequence (CCS) mode (PacBio, USA). For Hi-C sequencing, the cross-linked DNA was digested with Hind III enzyme, and paired-end (2 × 150 bp) sequenced on an Illumina NovaSeq 6000 system (Illumina, USA). For full-length transcript sequencing, we used RNeasy Plant Mini Kit (QIAGEN, Germany) to extract the total RNA from the root, stem, and leaf, respectively, and then pooled in equal amount for sequencing library preparation. The library with 0.5-6 kb cDNA fragments was sequenced with Iso-Seq mode on PacBio Sequel II system (PacBio, USA).

### Genome assembly

Before genome assembly, the low-quality PacBio HiFi reads that with read length < 2.000 bp or average quality < 99% were removed. The genome size was estimated by GCE v1.02 (https://github.com/fanagislab/GCE) based on K-mer frequencies using PacBio HiFi reads. Then, PacBio HiFi reads were assembled using Hifiasm v0.16.1 (Cheng *et al*., 2021) with parameter “-l 0”. The contaminated contigs were detected by aligning the contigs to the prokaryotic reference genomes and mitochondrion and plastid genomes using Minimap2 v2.20 (Li, 2018), and removed the contigs that have high identity (> 95%) and coverage (>95%) to the targets. The pseudo-chromosomes were constructed using Hi-C sequencing data. We used HiC-Pro v2.11.4 (Servant *et al*., 2015) pipeline to perform the Hi-C reads mapping, valid ligation pairs detection, and the Hi-C link matrixes generation, which were further used for pseudo-chromosomes construction by EndHiC v1.0 (Wang *et al*., 2022). The completeness of the genome assembly was estimated using BUSCO v5.2.2 (BUSCO) (Simao *et al*., 2015) against embryophyta_odb10 database.

### Genome annotation

For transposable element (TE) identification, we firstly used EDTA v1.9.9 (Ou *et al*., 2019) to produce a TE library based on the structural analysis. Secondly, the TE repeats were identified by homology-searching against the above structural TE library, protein-coding TE database, and Repbase v26.05 (Bao *et al*., 2015) using RepeatMasker v4.1.2 (Smit *et al*.). Thirdly, an extra *de novo* TE library was constructed from the non-TE sequences (masked all identified TEs with Ns) by RepeatModeler v2.0.2 (Flynn *et al*., 2020), and the library was used by RepeatMasker to identify the remaining TEs. Finally, we combined all identified TEs together. In addition, the tandem repeat (TR) elements were identified using Tandem Repeats Finder (TRF) v4.07 (Benson, 1999) with parameter “2 5 7 80 10 50 2000 -h -d”.

The protein-coding genes were prediction by using Augustus v3.4.0 (Stanke *et al*., 2006), with transcript and homology hints and parameter “--softmasking=on”. The transcript hints were obtained by mapping RNA sequences from PacBio Iso-Seq sequencing to the genome using GMAP v2020-10-27 (Wu and Watanabe, 2005), and the homology hints were generated by aligning the protein-coding sequences from the 12 published Papilionoideae species, including *Lotus japonicus, Medicago truncatula, Pisum sativum, Phaseolus vulgaris, Pueraria lobata* var. *montana, Lablab purpureus, Mucuna pruriens, Glycine max, Vigna angularis, Aeschynomene evenia, Arachis hypogaea*, and *Styphnolobium japonicum*, to the genome assembly by using Exonerate v2.2.0 (Slater and Birney, 2005). In addition, the tRNA genes were predicted using tRNAscan-SE v2.0 (Chan *et al*., 2021), and rRNA and other ncRNA genes were identified by cmscan from infernal v1.1.4 (Nawrocki and Eddy, 2013) based on Rfam v14 (Kalvari *et al*., 2021).

For gene functional annotation, the protein sequences of the genes were aligned to KEGG and NCBI-NR database by DIAMOND v2 (Buchfink *et al*., 2021), and the best-hit from the alignment results were chose for analyzing. In addition, the protein domain annotation was performed using InterProScan v5.52-86 (Blum *et al*., 2021) against InterPro database.

### Evolutionary analysis

The protein-coding sequences from yam bean, the 12 published Papilionoideae species, including *Lotus japonicus, Medicago truncatula, Pisum sativum, Phaseolus vulgaris, Pueraria lobata* var. *montana, Lablab purpureus, Mucuna pruriens, Glycine max, Vigna angularis, Aeschynomene evenia, Arachis hypogaea*, and *Styphnolobium japonicum*, and an outgroup species *Arabidopsis thaliana* were combined. Then, the orthogroups were constructed for these species by OrthoFinder v2.5.2 (Emms and Kelly, 2019) with parameter “-M msa -A mafft -T fasttree -l -y”, and a phylogenetic tree was constructed by STAG (https://github.com/davidemms/STAG) in OrthoFinder. The divergence time was investigated using MEGA v11 (Kumar *et al*., 2018), by setting the calibration time of 102.0 - 112.5 million years ago (MYA) between *Arabidopsis thaliana* and Fabaceae species, which was obtained from the website of TimeTree (www.timetree.org). In addition, to confirm the occurrence of the common legume whole-genome duplication (WGD) event, the collinear blocks of inter- and intra-species for the genome of yam bean and soybean were determined by MCScanX (Wang *et al*., 2012), and pairwise synonymous rates (*Ks*) of paralogous genes from collinear blocks were calculated by KaKs_calculator (Wang *et al*., 2010) with the GMYN model.

## Results

### Chromosome-level genome assembly of yam bean

We assembled the genome of yam bean (cv. Fanji) with combined datasets from PacBio HiFi and Hi-C sequencing. We generated 79.5 Gb (147 ×) PacBio HiFi read data with a read N50 size of 19.9 Kb, and assembled the data into contigs with Hifiasm (Cheng *et al*., 2021), followed by removing of the short contaminated contigs. The assembly includes 291 contigs with a N50 size of 25.6 Mb and total length of 539.0 Mb (Table 1), which was consistent with the estimated genome size (∼ 530 Mb) based on K-mer frequencies (Figure 1a). The assembled genome has a complete rate of 99.3% estimated using BUSCO (Simao *et al*., 2015) against the embryophyta lineage. Compared to the previously reported yam bean genome (Fernandez *et al*., 2021), the current genome assembly has a 1,388-fold increase in contig N50 size, and a 12.2% increase in BUSCO complete rate.

**Table 1.**
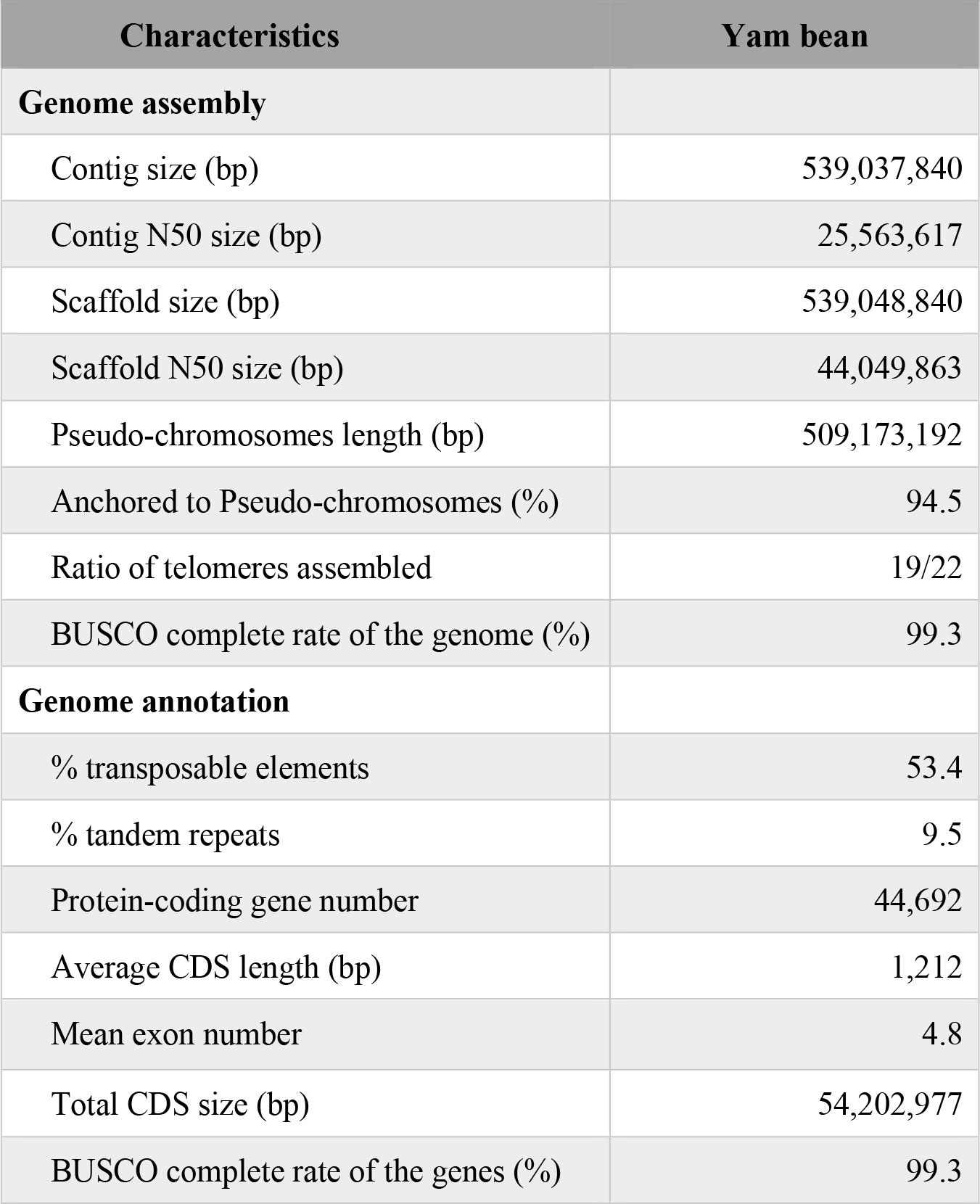
Statistics of the genome assembly and annotation for yam bean.

| Characteristics | Yam bean |
| --- | --- |
| **Genome assembly** |  |
| Contig size (bp) | 539,037,840 |
| Contig N50 size (bp) | 25,563,617 |
| Scaffold size (bp) | 539,048,840 |
| Scaffold N50 size (bp) | 44,049,863 |
| Pseudo-chromosomes length (bp) | 509,173,192 |
| Anchored to Pseudo-chromosomes (%) | 94.5 |
| Ratio of telomeres assembled | 19/22 |
| BUSCO complete rate of the genome (%) | 99.3 |
| **Genome annotation** |  |
| % transposable elements | 53.4 |
| % tandem repeats | 9.5 |
| Protein-coding gene number | 44,692 |
| Average CDS length (bp) | 1,212 |
| Mean exon number | 4.8 |
| Total CDS size (bp) | 54,202,977 |
| BUSCO complete rate of the genes (%) | 99.3 |

**Figure 1.**
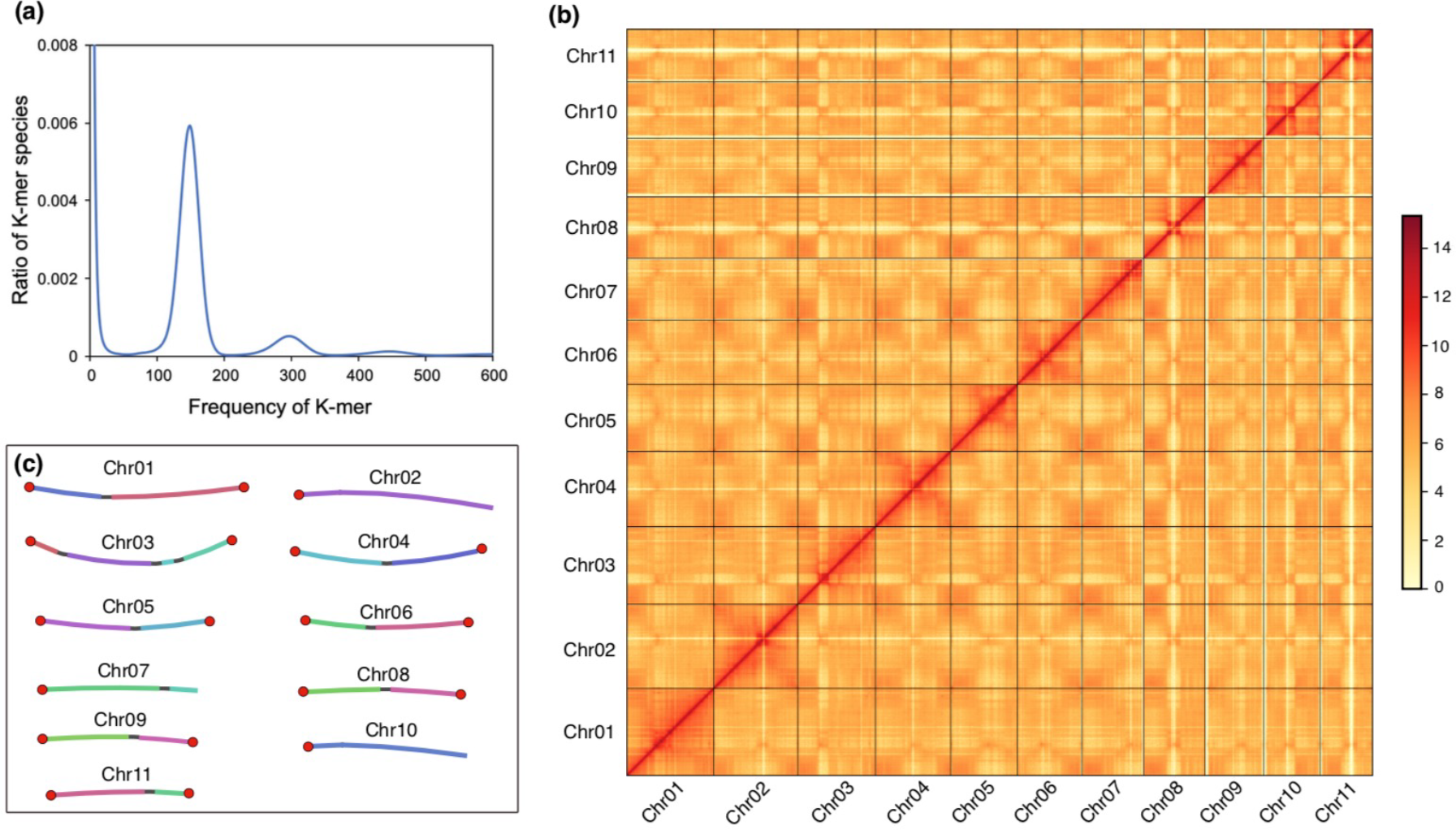
Assessment of the genome assembly for yam bean. (a) Distribution of K-mer frequencies in sequencing reads. The main peak reflects the “unique” regions in the genome. K-size equal 17; (b) Hi-C heatmap of the genome assembly. Bin size is 500-Kb, and color represents Log2(links number); (c) Bandage view of the pseudo-chromosomes. The colored lines represent contigs, and the gray lines represent the links between the two contigs. The chromosome ends assembled with telomeric repeats were highlighted with red solid circle.

Using 79.5 Gb (147 ×) of Hi-C data, we successfully anchored 509.2 Mb sequences (94.5% of the entire assembly) into 11 pseudo-chromosomes (Figure 1b), with 9 contains two contigs and 2 contains only one contig (Figure 1c). Then, the telomeric repeats of the pseudo-chromosomes were analyzed based on the result from TRF (Benson, 1999), revealing that 86.4% (19/22) of the assembled chromosome-ends have telomeric repeats (Table 1), with 8 pseudo-chromosomes have the repeats at the both ends (Figure 1c). In summary, we assembled a nearly complete high-quality chromosome-level reference genome for yam bean with the N50 and N90 size of 44.0 and 35.3 Mb, and 8 of the 11 pseudo-chromosomes have telomeres at the both ends.

### High-quality gene set of yam bean

To analyze the transposable elements (TE), we searched the genome sequence by using a combination of *de novo* and homology-based strategies, and found that 53.4% (287.7 Mb) of yam bean genome consisted of TEs (Table 1), which was similar to that of the closely related species soybean (∼ 54%) (Liu *et al*., 2020). Among the annotated TEs, we identified 2,380 intact TEs (with total length of 17.2 Mb) that with clear structural boundaries. Long terminal repeats (LTR) is the most abundant component of TEs, accounting for 28.8% of the genome (Table 2), with Gypsy (21.9%) and Copia (10.7%) as the major members. In addition, we investigated the tandem repeats (TR) using TRF (Benson, 1999), and found 51.0 Mb (9.5% of the genome) of TRs in yam beam genome (Table 1).

**Table 2.** Statistics of transposable element content in various classes.

| TE class | Length (bp) | % of genome |
| --- | --- | --- |
| LTR | 155,240,188 | 28.80 |
| DNA elements | 111,299,756 | 20.65 |
| LINE | 14,776,314 | 2.74 |
| SINE | 2,645,274 | 0.49 |
| MITE | 1,532,778 | 0.28 |
| RC | 390,527 | 0.07 |
| Other | 1,767,419 | 0.33 |
| Total | 287,652,256 | 53.36 |

Note: LTR, long terminal repeat; LINE, long interspersed nuclear element; SINE, short interspersed element; MITE, miniature inverted-repeat transposable element; RC, rolling-circle transposable element.

A total of 44,692 protein-coding genes were predicted by using Augustus (Stanke *et al*., 2006), with an average coding sequence (CDS) length of 1,212 bp and a mean exon number of 4.8 (Table 1). The BUSCO complete rate of the genes were 99.3%, which was consistent with that of the assembled genome sequences (99.3%) (Table 1), suggesting the high completeness of the genes and the assembled genome. In addition, the BUSCO complete rate was obviously higher than that of the previously reported (75%) (Fernandez *et al*., 2021). Functional annotation of the predicted genes indicated that 88.7% of genes were annotated by NCBI-NR, KEGG, and InterPro database. In addition, we identified 1,970 tRNA genes and 5,766 rRNA genes (5S 1,431, 5.8S 1,365, 18S 1,475, and 28S 1,495) in the assembled genome, and 1,608 of other ncRNA genes.

### Evolutionary analysis

To investigate the divergence time of yam bean and the closely related species, we firstly combined the reference genes from yam bean, 12 published Papilionoideae species (*Lotus japonicus, Medicago truncatula, Pisum sativum, Phaseolus vulgaris, Pueraria lobata* var. *montana, Lablab purpureus, Mucuna pruriens, Glycine max, Vigna angularis, Aeschynomene evenia, Arachis hypogaea*, and *Styphnolobium japonicum*), and an outgroup species *Arabidopsis thaliana*. Then, these genes were clustered into 31,542 orthogroups by using OrthoFinder (Emms and Kelly, 2019), and a species tree was built based on 1,246 orthogroups that with minimum of 86.7% of species having single-copy genes in any orthogroup. The topology of the resulted phylogenetic tree showed that yam bean was closely related to soybean (*Glycine max*) and *Pueraria lobata* var. *montana* (Figure 2a), which was consistent with the result from a previous study (Fernandez *et al*., 2021).

**Figure 2.**
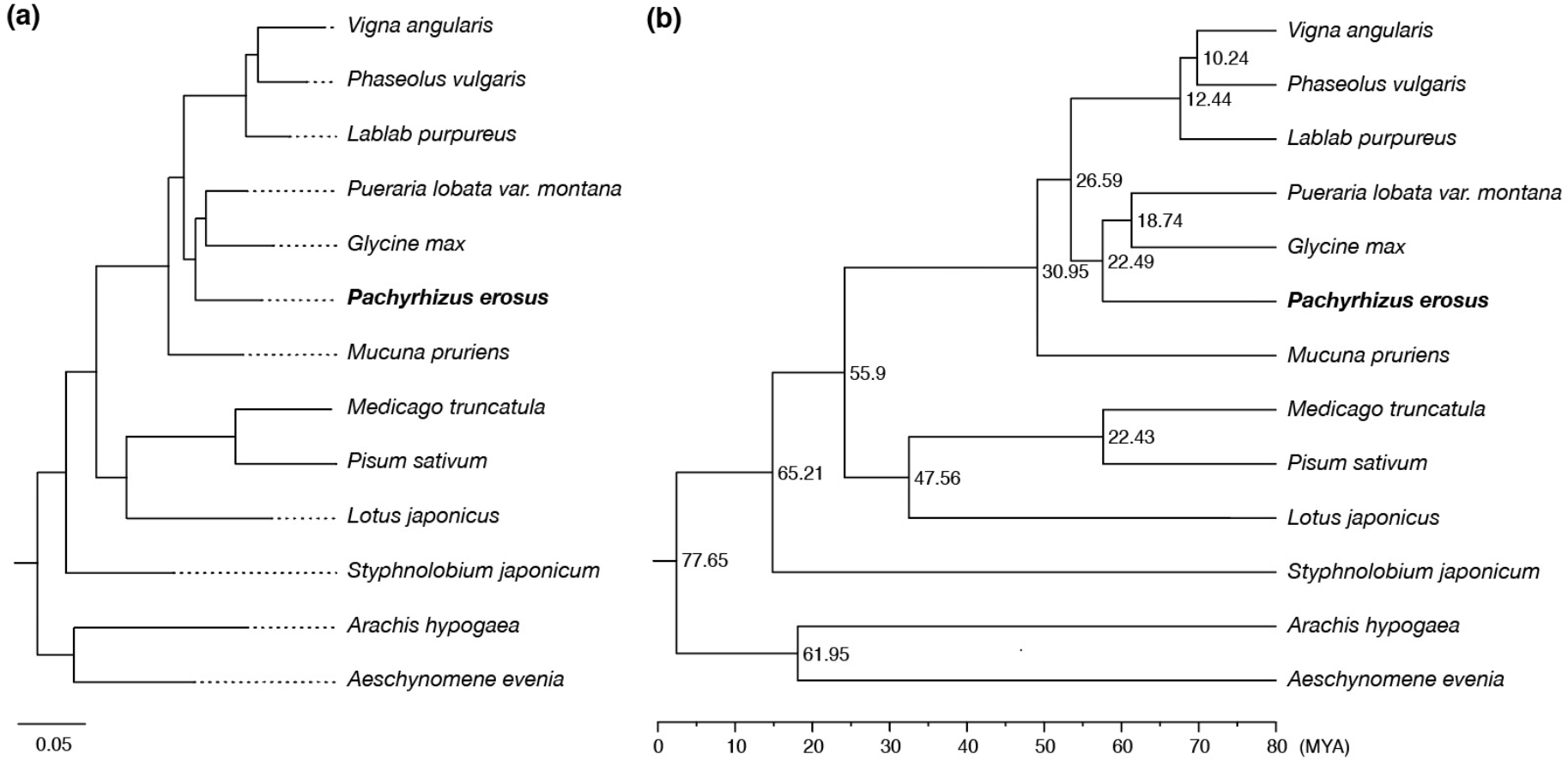
Phylogenetic analysis for yam bean. (a) Phylogeny tree constructed by STAT in OrthoFinder. The outgroup species of *Arabidopsis thaliana* was not shown. The bar means substitution per amino acid site; (b) The divergence time was estimated by MEGA, and setting the calibration time of 102.0 - 112.5 million years ago (MYA) between *Arabidopsis thaliana* and Fabaceae species. MYA, million years ago. The node labels indicate estimated divergence time.

The divergence time on the phylogenetic tree was estimated by MEGA (Kumar *et al*., 2018), using the divergence time between the species *Arabidopsis thaliana* and the family *Fabaceae* (102.0 - 112.5 MYA) as a calibration constraint. The result showed that soybean and *Pueraria lobata* var. *montana* diverged from each other 18.7 MYA (Figure 2b), which was consistent with a previous reported 18.2 MYA (Mo *et al*., 2022), revealing the correctness of our divergence time estimation. As a result, we found that yam bean diverged from the clade of soybean and *Pueraria lobata* var. *montana* 22.5 MYA (Figure 2b). In addition, we identified 12,190 (27.3%) intra-species syntenic genes within yam bean by MCScanX (Wang *et al*., 2012), and then the *Ks* values of the paralog pairs in the syntenic fragments were calculated. The *Ks* distributions of yam bean showed an obvious peak at ∼ 0.5 (Figure 3), which was consistent with that of soybean (Schmutz *et al*., 2010). Besides, the peak occurred before the speciation peak at 0.15 (Figure 3), confirming the occurrence of the common legume whole-genome duplication (WGD) event (Schmutz *et al*., 2010).

**Figure 3.**
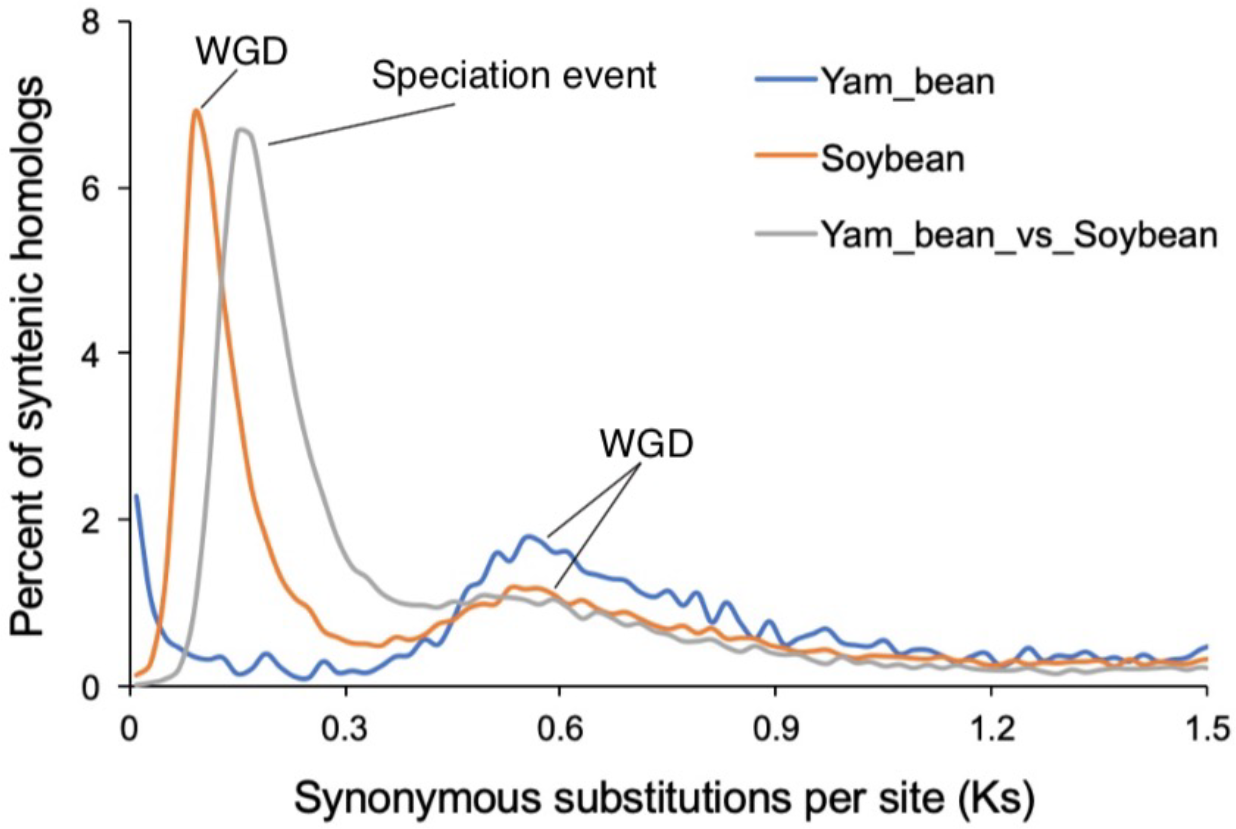
Ks distribution of orthologous or paralogous genes for yam bean and soybean

## Discussion

In this study, we generated a high-quality chromosome-level genome assembly for yam bean based on the highly accurate PacBio HiFi long reads and Hi-C sequencing data. Compared to the previously reported yam bean genome (Fernandez *et al*., 2021), the current assembly has a 1,388-fold increase in contig N50 size, and 12.2% and 24.3% increase in BUSCO complete rate for the genome sequence and gene set, respectively, suggesting an obviously high improvement of our genome assembly. Evolutionary analysis revealed that yam bean diverged from the clade of soybean and *Pueraria lobata* var. *montana* 22.5 MYA, and the occurrence of the common legume whole-genome duplication (WGD) event were confirmed based on *Ks* analysis. This high-quality genome assembly will greatly facilitate the breeding of yam bean based on the genetic and genomic methods.

## Data Availability Statement

All the data that generated during the current study have been deposited at DDBJ/ENA/GenBank under project accession PRJNA1002465. The genome assembly and annotations are available at Zenodo: https://www.zenodo.org/record/8372125.

## Literature Cited

Adewale BD, Nnamani CV. Introduction to food, feed, and health wealth in African yam bean, a locked-in African indigenous tuberous legume. Frontiers in Sustainable Food Systems. 2022;6.

Bao W, Kojima KK, Kohany O. Repbase Update, a database of repetitive elements in eukaryotic genomes. Mob DNA. 2015;611.

Benson G. Tandem repeats finder: a program to analyze DNA sequences. Nucleic Acids Res. 1999;27(2):573–580.

Blum M, Chang HY, Chuguransky S, Grego T, Kandasaamy S, Mitchell A et al. The InterPro protein families and domains database: 20 years on. Nucleic Acids Res. 2021;49(D1):D344–D354.

Buchfink B, Reuter K, Drost HG. Sensitive protein alignments at tree-of-life scale using DIAMOND. Nat Methods. 2021;18(4):366.

Chan PP, Lin BY, Mak AJ, Lowe TM. tRNAscan-SE 2.0: improved detection and functional classification of transfer RNA genes. Nucleic Acids Res. 2021;49(16):9077–9096.

Cheng HY, Concepcion GT, Feng XW, Zhang HW, Li H. Haplotype-resolved de novo assembly using phased assembly graphs with hifiasm. Nat Methods. 2021;18(2):170.

Council NR (2006). Lost Crops of Africa: Volume II: Vegetables. The National Academies Press: Washington, DC.

Emms DM, Kelly S. OrthoFinder: phylogenetic orthology inference for comparative genomics. Genome Biol. 2019;20(1):238.

Fernandez CGT, Pati K, Severn-Ellis AA, Batley J, Edwards D. Studying the Genetic Diversity of Yam Bean Using a New Draft Genome Assembly. Agronomy-Basel. 2021;11(5).

Flynn JM, Hubley R, Goubert C, Rosen J, Clark AG, Feschotte C, Smit AF. RepeatModeler2 for automated genomic discovery of transposable element families. Proc Natl Acad Sci U S A. 2020;117(17):9451–9457.

Kalvari I, Nawrocki EP, Ontiveros-Palacios N, Argasinska J, Lamkiewicz K, Marz M et al. Rfam 14: expanded coverage of metagenomic, viral and microRNA families. Nucleic Acids Res. 2021;49(D1):D192–D200.

Kumar S, Stecher G, Li M, Knyaz C, Tamura K. MEGA X: Molecular Evolutionary Genetics Analysis across Computing Platforms. Mol Biol Evol. 2018;35(6):1547–1549.

Li H. Minimap2: pairwise alignment for nucleotide sequences. Bioinformatics. 2018;34(18):3094–3100.

Liu Y, Du H, Li P, Shen Y, Peng H, Liu S et al. Pan-Genome of Wild and Cultivated Soybeans. Cell. 2020;182(1):162–176 e113.

Mo CJ, Wu ZD, Shang XH, Shi PL, Wei MH, Wang HY et al. Chromosome-level and graphic genomes provide insights into metabolism of bioactive metabolites and cold-adaption of Pueraria lobata var. montana. DNA Research. 2022;29(5).

Nawrocki EP, Eddy SR. Infernal 1.1: 100-fold faster RNA homology searches. Bioinformatics. 2013;29(22):2933–2935.

Ou S, Su W, Liao Y, Chougule K, Agda JRA, Hellinga AJ et al. Benchmarking transposable element annotation methods for creation of a streamlined, comprehensive pipeline. Genome Biol. 2019;20(1):275.

Schmutz J, Cannon SB, Schlueter J, Ma J, Mitros T, Nelson W et al. Genome sequence of the palaeopolyploid soybean. Nature. 2010;463(7278):178–183.

Servant N, Varoquaux N, Lajoie BR, Viara E, Chen CJ, Vert JP, Heard E, Dekker J, Barillot E. HiC-Pro: an optimized and flexible pipeline for Hi-C data processing. Genome Biol. 2015;16259.

Simao FA, Waterhouse RM, Ioannidis P, Kriventseva EV, Zdobnov EM. BUSCO: assessing genome assembly and annotation completeness with single-copy orthologs. Bioinformatics. 2015;31(19):3210–3212.

Slater GS, Birney E. Automated generation of heuristics for biological sequence comparison. BMC Bioinformatics. 2005;631.

Smit A, Hubley R, Green P. RepeatMasker Open-4.0. 2015. http://repeatmasker.org/. Accessed: 10 November, 2020.

Stanke M, Schoffmann O, Morgenstern B, Waack S. Gene prediction in eukaryotes with a generalized hidden Markov model that uses hints from external sources. BMC Bioinformatics. 2006;7(1):62.

Wang D, Zhang Y, Zhang Z, Zhu J, Yu J. KaKs_Calculator 2.0: a toolkit incorporating gamma-series methods and sliding window strategies. Genom Proteom Bioinf. 2010;8(1):77–80.

Wang S, Wang HC, Jiang F, Wang AQ, Liu HW, Zhao HB, Yang BY, Xu D, Zhang Y, Fan W. EndHiC: assemble large contigs into chromosome-level scaffolds using the Hi-C links from contig ends. BMC Bioinformatics. 2022;23(1):528.

Wang YP, Tang HB, DeBarry JD, Tan X, Li JP, Wang XY et al. MCScanX: a toolkit for detection and evolutionary analysis of gene synteny and collinearity. Nucleic Acids Res. 2012;40(7):e49.

Wu TD, Watanabe CK. GMAP: a genomic mapping and alignment program for mRNA and EST sequences. Bioinformatics. 2005;21(9):1859–1875.

